# Excess extracellular K^+^ causes inner hair cell ribbon synapse degeneration

**DOI:** 10.1101/2019.12.23.887596

**Authors:** Hong-Bo Zhao, Yan Zhu, Li-Man Liu

## Abstract

Inner hair cell (IHC) ribbon synapses are the first synapse in the auditory system and can be degenerated by noise and aging, thereby leading to hidden hearing loss (HHL) and other hearing disorders. However, the mechanism underlying this cochlear synaptopathy remains unclear. Here, we report that elevation of extracellular K^+^, which is a consequence of noise exposure, could cause IHC ribbon synapse degeneration and swelling. Like intensity dependence in noise-induced cochlear synaptopathy, the K^+^-induced degeneration was dose-dependent, and could be attenuated by BK channel blockers. However, application of glutamate receptor (GluR) agonists caused ribbon swelling but not degeneration. In addition, consistent with synaptopathy in HHL, both K^+^ and noise exposure only caused IHC but not outer hair cell ribbon synapse degeneration. These data reveal that K^+^ excitotoxicity can degenerate IHC ribbon synapses in HHL, and suggest that BK channel may be a potential target for prevention and treatment of HHL.

## Introduction

Noise is a common factor for hearing loss. It has been found that exposure to moderate levels of noise, even short-term, can cause extensive degeneration of ribbon synapses between inner hair cells (IHCs) and auditory nerves without hair cell loss (*1-4*). This cochlear synaptopathy can cause hidden hearing loss (HHL), which presents with a normal audiogram with difficulty in hearing speech in noisy environments (*2, 5-6*). The cochlear IHC synapse degeneration also could impair hearing function in many aspects, such as neural adaptation, sound location, and temporal and speech processing (*7-9*). Such complaints account for ∼10% of patients who present to otolaryngology clinics (*7,10*). In addition, the ribbon synapse degeneration is observed at the early stage of age-related hearing loss and may accelerate its progression (*2, 11-14*). However, the underlying mechanism for the cochlear synapse degeneration remains unclear.

The ribbon synapses between IHCs and auditory nerves are composed of presynaptic ribbons in IHCs and postsynaptic glutamate receptors (GluR) at endings of auditory nerves. Glutamate is a major neurotransmitter of IHC ribbon synapses (*15*). It has been hypothesized that noise exposure may increase glutamate release, thereby leading to glutamate excitotoxicity which then causes hair cell death and synapse degeneration (*16-18*). However, HHL shows ribbon synapse degeneration without any hair cell loss. Also, the glutamate excitotoxicity mainly occurs at the postsynaptic GluRs located at the auditory nerve ending. Thus, it is hard to explain severe presynaptic ribbon degeneration in hair cells. Moreover, it was found that the presynaptic and postsynaptic components remained intact, despite the substantial terminal swelling following perfusion of GluR agonists (*19,20*). In addition, a recent study demonstrated that noise could still induce ribbon synapse swelling in mice lacking vesicular glutamate transporter-3 with deficiency in release of glutamate (*18*). These data suggest that other mechanisms then glutamate excitotoxicity have an important role in the ribbon synapse degeneration.

On the other hand, noise-induced hair cell and auditory nerve excessive-excitation increases K^+^ efflux from hair cells and nerve fibers, which can elevate extracellular (perilymphatic) K^+^ concentration causing K^+^ excitotoxicity. Noise can also impair the reticular lamina integrity breaking the barrier between endolymph and perilymph, thereby leading to K^+^-rich endolymph (K^+^ =∼150 mM) entering into the low-K^+^ perilymph (∼5 mM K^+^) causing K^+^ toxicity (*21, 22*). In the brain, it has been found that elevation of extracellular K^+^ resulting from cell death in the stroke area can cause secondary neuron degeneration in the neighboring non-stroke area (*23, 24*). In this study, we tested whether high extracellular K^+^ could cause cochlear synapse degeneration. We found that elevation of extracellular K^+^ concentration could cause IHC ribbon synapse degeneration and swelling in a dose-dependent manner. Blockage of K^+^ channels attenuated this degeneration. However, application of GluR agonists did not cause IHC ribbon synapse degeneration, even though they caused ribbon swelling. These data suggest that K^+^ excitotoxicity may play a major role in the IHC ribbon synapse degeneration, thereby shedding light on the prevention and treatment of this common type of hearing loss.

## Results

### Degeneration of IHC ribbon synapses by elevation of extracellular K^+^

To test whether high extracellular K^+^ could cause IHC ribbon synapse degeneration, we incubated one cochlea in the high-K^+^ extracellular solution which contained 50 mM K^+^ for 2 hr, as the same time as noise exposure to induce HHL. Another cochlea of the same mouse was simultaneously incubated in normal extracellular solution (NES) as control. Figure 1 shows that the IHC ribbon synapses were significantly reduced after challenge with 50 mM K^+^. In comparison with those in the NES group, the pre-synaptic ribbon (labeling puncta for CtBP2), post-synaptic GluR, and synapse (overlapped labeling) per IHC in the high-K^+^ group were reduced to 62.5±3.20%, 83.4±5.94%, and 60.8±5.58% (P=1.4E-05, 0.02, and 0.0008, paired t test, two-tail), respectively (Figure 1b). The reductions in ribbons and synapses were significantly larger than that in the GluR (P=0.022 and 0.013, respectively, one-way ANOVA with a Bonferroni correction). However, there was no significant difference between ribbon reduction and synapse reduction (P=1.00, one-way ANOVA). The percentages of synaptic ribbons and free ribbons in the high-K^+^ group and the NES group were almost the same (Supplementary Figure 1a&b) and had no significant difference between two groups (P=0.68, paired t test, two-tail), indicating that high-K^+^ caused both synaptic ribbon and free ribbon degeneration similarly. Thus, in the following analyses, we would not distinguish synaptic ribbon and free ribbon degeneration and analyze the total ribbon degeneration. Figure 1c shows that the total ribbons in the apical, middle, and basal cochlear turn in the high-K^+^ group were significantly reduced to 72.1±5.80%, 56.8±3.73%, and 57.2±7.65%, respectively, in comparison with those in the control group with NES incubation (P=0.006, 5.4E-06, and 0.004, paired t test, two-tail). The reduction in the apical turn appeared less than that in the middle and basal cochlear turns. However, there was no significant difference among them (P=0.13, one-way ANOVA). High-K^+^ also caused ribbon swelling (Figure 1d). In comparison with that in the control group with NES incubation, the ribbon volume in the high-K^+^ group was significantly increased 1.80±0.22 folds (P=0.003, paired t test, two-tail).

**Fig 1.**
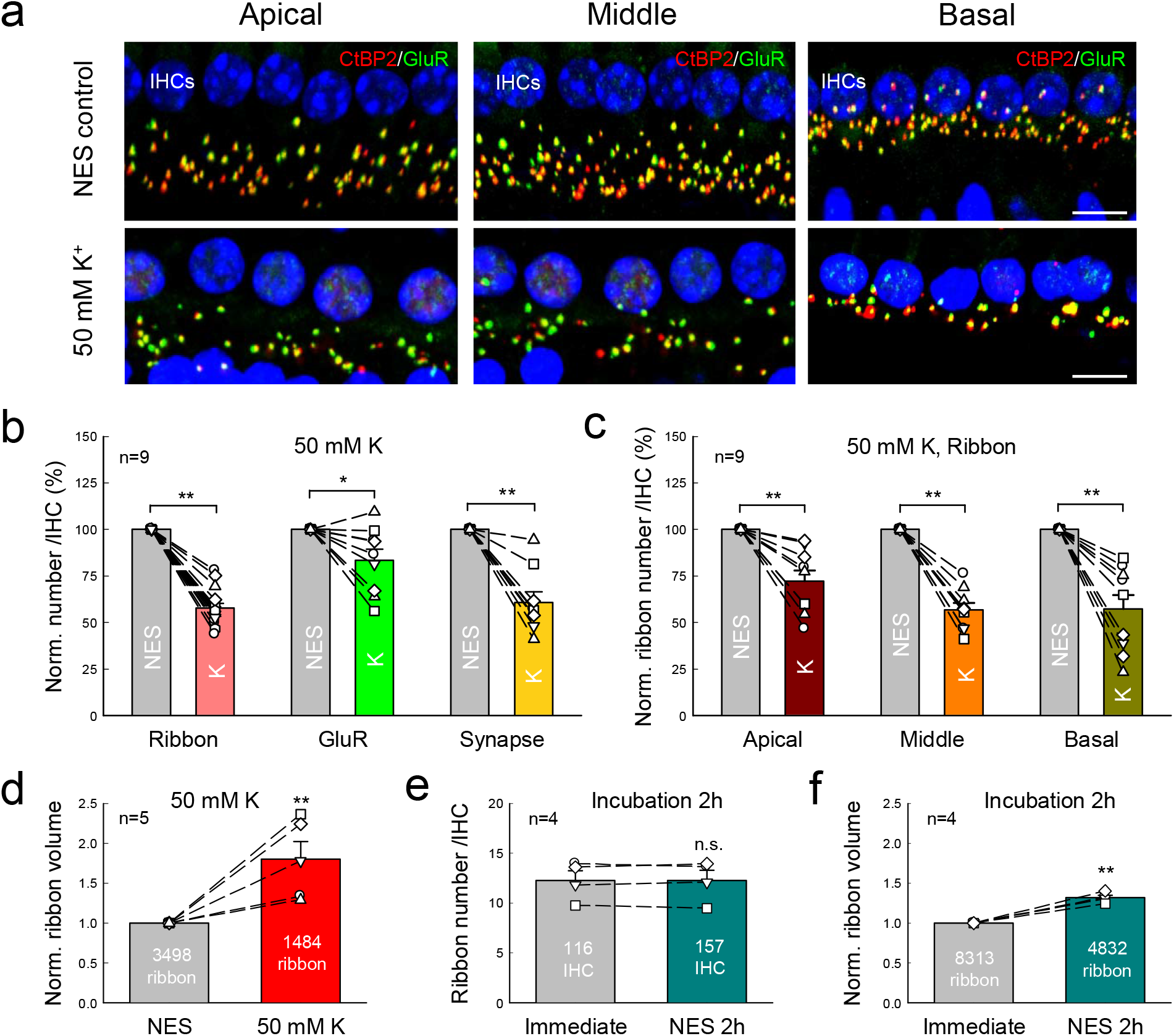
High-K^+^ induced IHC ribbon synapse degeneration. **a:** Immunofluorescent staining for ribbon synapses at the IHC area in the apical, middle, and basal turns after incubation. Two cochleas of a mouse were incubated separately in the high-K^+^ (50 mM) extracellular solution and in the normal extracellular solution (NES) for 2 hr. Presynaptic ribbons are labeled by CtBP2 (red) and postsynaptic GluRs are labeled by green. Scale bar: 10 µm. **b:** Quantitative analyses of high-K^+^ induced ribbon synapse degeneration at IHCs. Pre-synaptic ribbons, post-synaptic GluRs, and synapses per IHC pooled from all cochlear turns were averaged and normalized to those in the control cochlea of the same mouse with NES incubation. Dashed lines connected two same symbols represent data collected from two cochleas of the same mouse. N is the animal number. **c:** K^+^-induced ribbon degeneration in the cochlear apical, middle, and basal turn. The ribbon numbers in the different turns were normalized to those in the corresponding turn in the control cochlea of the same mouse. **d:** High-K^+^ induced ribbon swelling. The volume of labeling puncta for CtBP2 were measured. The data pooled from all cochlear turns were averaged and normalized to that in the control cochlea of the same mouse. Total numbers of measured ribbons are presented within the bars. N is the animal number. **e-f:** Incubation procedure has no effect on ribbon degeneration. One cochlea was immediately fixed after isolation and stained for CtBP2 and another cochlea of the same mouse was incubated in NES for 2 hr and then fixed for staining. Dashed lines connected two same symbols represent data collected from two cochleas of the same mouse. Data were pooled from all cochlear turns. Numbers in the bars represent the total numbers of accounted IHCs and measured ribbons. N is the animal number. *: P<0.05, **: P<0.01, paired t test, two-tail for all panels.

We also tested the effect of the incubation procedure on the ribbon degeneration (Figure 1e&f). One cochlea was incubated in the NES for 2 hr and another cochlea in the same mouse was immediately fixed without incubation after isolation. The ribbon numbers in the incubation group and non-incubation group were 12.3±1.01/IHC and 12.3±0.96/IHC, respectively (Figure 1e). There was no significant difference between them (P=0.98, paired t test, two-tail). The ribbon volume after incubation had a small increase and was increased 1.32 folds, i.e., increase of 32% (Figure 1f, P=0.002, paired t test, two-tail). However, in comparison with the increase of 2.371 folds (i.e., increase of 137.1%, re: non-incubation group) in the high-K^+^ group (Supplementary Figure 3b), the volume increase caused by the incubation procedure was significantly small (P=0.02, one-way ANOVA with a Bonferroni correction).

K^+^-induced ribbon degeneration is dose-dependent (Figure 2). As extracellular K^+^ concentration increased, the ribbon degeneration was increased. When the extracellular K^+^-concentration was greater than 30 mM, the ribbon degeneration reached a stable stage level. The median concentration of K^+^ was at 24.7 mM.

**Fig 2.**
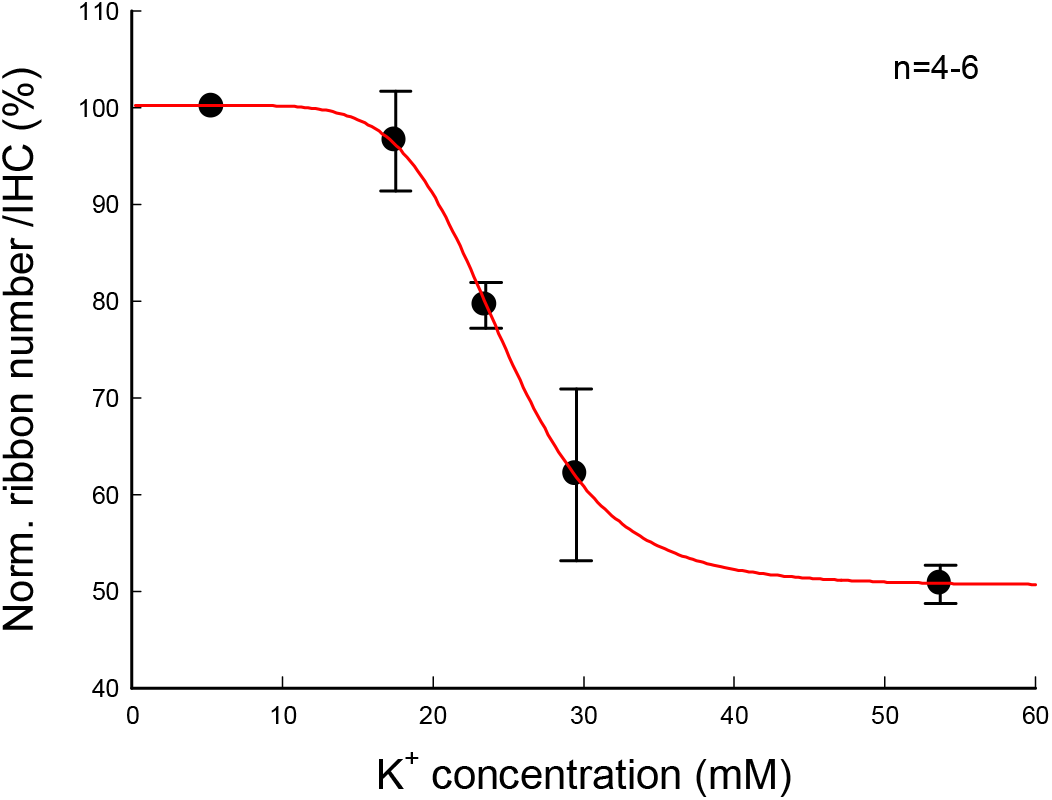
Dose-curve of K^+^-induced ribbon degeneration in IHCs. One cochlea was incubated with one concentration of K^+^ and another cochlea of the same mouse was incubated in the NES (K^+^=5.37 mM) as control. The data pooled from all cochlear turns were averaged and normalized to that in the control cochlea of the same mouse. N is the mouse number at each data point. A smooth solid line represents data fitting to a Hill’s function *y=y*_*0*_*+a*C*^*n*^*/(EC*_*50*_ ^*n*^*+C*^*n*^), where median concentration *EC*_*50*_ =24.7 mM.

### Attenuation of IHC ribbon synapse degeneration by K^+^ channel blockers

We further tested whether K^+^ channel blockers could attenuate this high-K^+^ induced IHC ribbon synapse degeneration. Since the Ca^++^-dependent BK channel is a prevalent K^+^ channel at IHCs (*25, 26*), the specific BK channel blocker iberiotoxin (IBT) was used (Figure 3a&b). We incubated one cochlea in 50 mM K^+^ extracellular solution with 0.1 μM IBT (K^+^ +IBT group) and another cochlea of the same mouse in the 50 mM K^+^ extracellular solution (high-K^+^ group) as control. In comparison with those in the high-K^+^ control group, the survival of ribbons, GluRs, and synapses in the K^+^+IBT group were increased by 151.5±8.36%, 128.9±7.73%, and 153.4±14.0% (P=0.0035, 0.0201, and 0.0189, paired t test, two-tail), respectively (Figure 3a). In the apical, middle, and basal turn, the survival of ribbons was increased by 154.3±9.89%, 151.7±13.5%, and 165.5±14.0% (P=0.0029, 0.0093, and 0.0094, paired t test, two-tail), respectively (Figure 3b). If compared with those in the NES control group (Figure 1c), the ribbon survival in the apical, middle, and basal turns in the K^+^+IBT group was increased to 78.8±3.15, 78.0±2.12, and 81.6±11.8%, respectively, for 50 mM K^+^ challenge. However, there was no significant difference in the increments among different cochlear turns (P=0.72, one-way ANOVA). Thus, IBT could significantly increase IHC ribbon synapse survival for high-K^+^ challenge.

**Fig 3.**
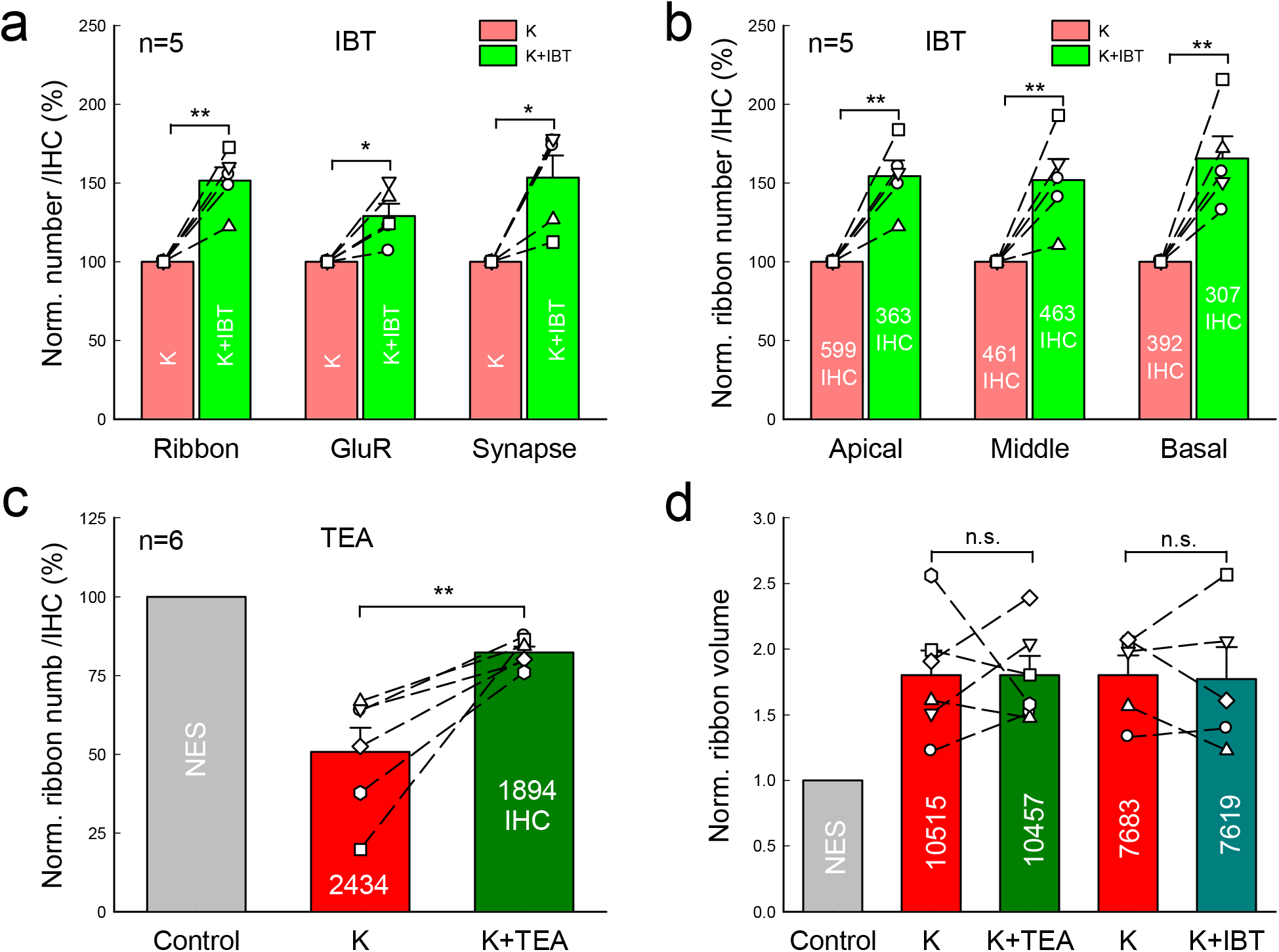
Attenuation of IHC ribbon synapse degeneration by K^+^ channel blockers. **a:** Attenuation of the BK channel blocker iberiotoxin (IBT) on K^+^-induced IHC ribbon synapse degeneration. One cochlea was incubated in 50 mM K^+^ extracellular solution (high-K^+^ group) and another cochlea of the same mouse was incubated in 50 mM K^+^ extracellular solution with 0.1 μM IBT (K^+^+IBT group). Both cochleas were simultaneously incubated for 2 hr. The numbers of presynaptic ribbons, postsynaptic GluRs, and synapses per IHC from all cochlear turns were averaged and normalized to those in the high-K^+^ group. Dashed lines connected two same symbols represent data collected from two cochleas of the same mouse. N is the mouse number. **b:** Attenuation of ribbon degeneration by IBT in different cochlear turns. Total numbers of accounted IHCs are presented in each bar. **c:** Attenuation of a K^+^ channel blocker TEA on ribbon degeneration. Two cochleas of a mouse were incubated separately in the high-K^+^ (50 mM) extracellular solution (high-K^+^ group) and in the 50 mM K^+^ extracellular solution with 20 mM TEA (K^+^+TEA group) for 2 hr. Ribbon numbers per IHC from all cochlear turns in the high-K^+^ group and the K^+^+TEA group were separately calculated and normalized to that in the NES control group in Figure 1. Dashed lines connected two same symbols represent data collected from two cochleas of the same mouse. **d:** IBT and TEA have no significant attenuating effect on K^+^-induced ribbon swelling. Ribbon volumes were pooled from all cochlear turns, and averaged and normalized to that in the NES control group in Figure 1. Numbers in the bars represent the total numbers of measured ribbons. *: P < 0.05, **: P<0.01, n.s.: no significance (P>0.05), paired t test, two-tail for all panels.

Since IHCs also have other K^+^ channels (*27, 28*), we further used a broad K^+^ channel blocker tetraethylammonium (TEA). As the same as IBT, one cochlea was incubated with 50 mM K^+^ extracellular solution with 20 mM TEA (K^+^+TEA group) and another cochlea of the same mouse was incubated in 50 mM K^+^ extracellular solution (high-K^+^ group). Both ribbon numbers in the K^+^-TEA group and the high-K^+^ group were normalized to that in the NES control group in Figure 1. Figure 3c shows that 82.2±1.85% of IHC ribbons in the K^+^+TEA group was reserved for 50 mM K^+^ challenge (P=0.01, paired t test, two-tail). In comparison with that (79.5±6.47%) in the K^+^+IBT group, there was no significant difference between them (P=0.70, t test, two-tail). However, in comparison with that in the NES control group in Figure 1, the ribbon volumes in the K^+^+IBT group and the K^+^+TEA group were still increased 1.77±0.24 folds and 1.80±0.15 folds (Figure 3d), respectively, and had no significant differences from those in the high-K^+^ control groups (P=0.88 and 0.95, respectively, paired t test, two-tail). Thus, both IBT and TEA had no effect on the ribbon swelling.

### Attenuation of Ca^++^ channel blockers on IHC ribbon synapse degeneration and swelling

Since BK channels could be Ca^++^-dependent, we further tested whether blockage of Ca^++^ channels could attenuate the high-K^+^ induced IHC ribbon synapse degeneration. Co^++^ is a broad Ca^++^ channel blocker. As the same as application of IBT and TEA, we added 2 mM Co^++^ into the high-K^+^ (50 mM) incubation solution for one cochlea incubation (K^+^+Co^++^ group) and another cochlea of the same mouse was incubated with 50 mM K^+^ extracellular solution (high-K^+^ group). The ribbon numbers in both groups were normalized to that in the NES control group in Figure 1. Figure 4a showed that the ribbon survival was significantly increased from 50.8±7.00% in the high-K^+^ group to 72.7±7.07% in the K^+^+Co^++^ group (P=0.006, paired t test, two-tail). The ribbon swelling (Figure 4b) was also significantly reduced from 1.80±0.16 folds in the high-K^+^ group to 1.29±0.14 folds in the K^+^+Co^++^ group (P=0.005, paired t test, two-tail). Thus, the Ca^++^ channel blocker Co^++^ could attenuate both ribbon degeneration and swelling.

**Fig 4.**
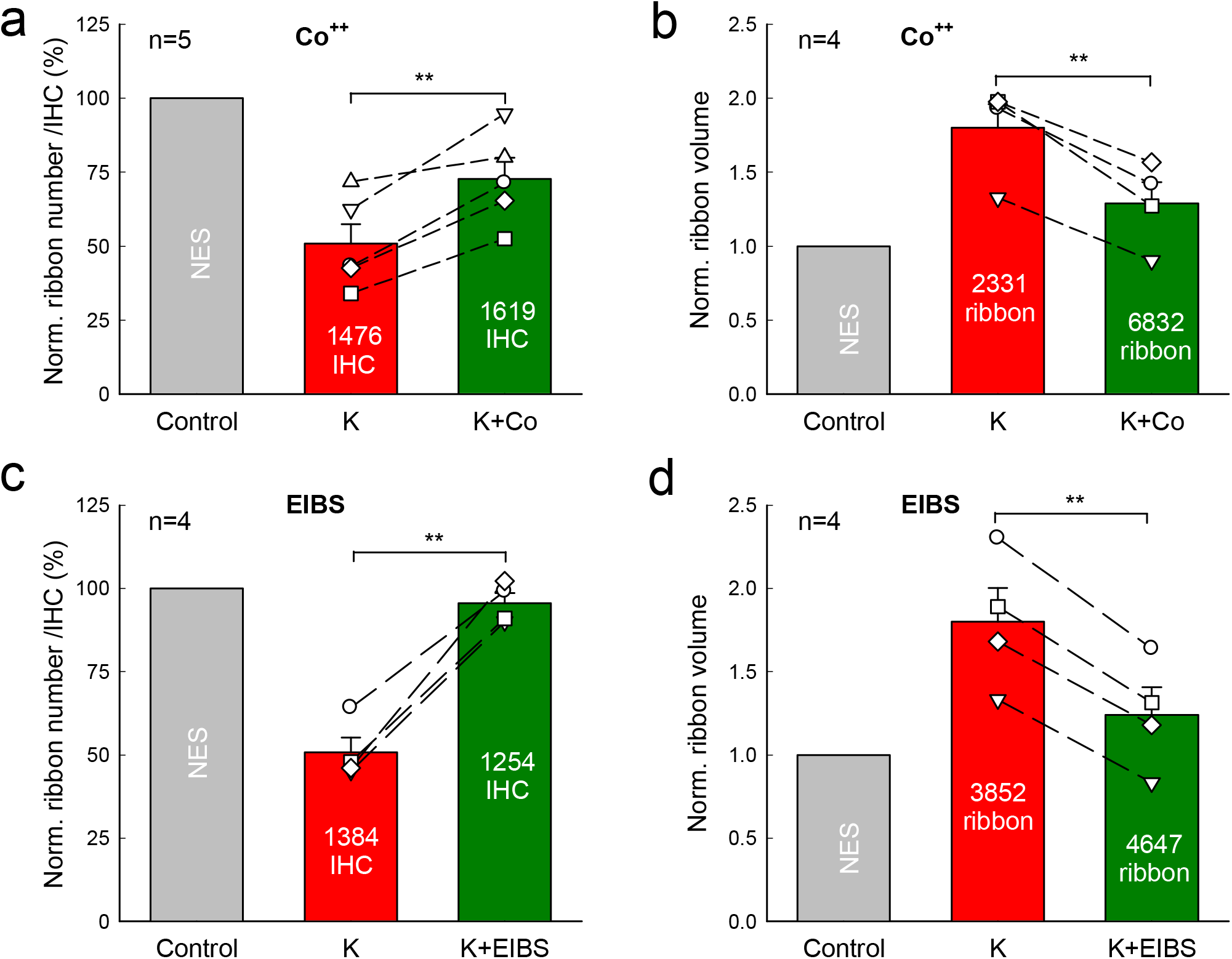
Attenuation of IHC ribbon degeneration and swelling by Ca^++^ channel blocker. **a-b:** Attenuation of the Ca^++^ channel blocker Co^++^ on ribbon degeneration and swelling induced by high-K^+^ challenge. Two cochleas of the same mouse were incubated separately in the 50 mM K^+^ extracellular solution with 2 mM Co^++^ (K^+^+Co^++^ group) and 50 mM K^+^ extracellular solution (high-K^+^ group). Ribbon numbers and volumes pooled from all cochlear turns in both groups were separately averaged and normalized to that in the NES group in Figure 1. Dashed lines connected two same symbols represent data collected from two cochleas of the same mouse. N is the animal number. Number in each bar represents accounted IHC or ribbon numbers. Co^++^ significantly attenuates both ribbon degeneration and swelling induced by high-K^+^. **c-d:** Attenuation of high-K^+^ induced ribbon degeneration and swelling by extracellular ionic blocking solution (EIBS), which contains 20 mM TEA, 20 mM Cs^+^, and 2 mM Co^++^ and can block both Ca^++^ and K^+^ channels extensively in the hair cells. One cochlea in a mouse was incubated in the high-K^+^ (50 mM) extracellular solution (high-K^+^ group) and another cochlea was incubated in the 50 mM K^+^ with EIBS (K^+^+EIBS group) for 2 hr. Ribbon numbers and volumes in both the high-K^+^ group and the K^+^+EIBS group were separately normalized to those in the NES control group in Figure 1. **: P < 0.01, paired t test, two-tail.

We also used the extracellular ionic blocking solution (EIBS), which contains 20 mM TEA, 2 mM Co^++^, and 20 mM Cs^+^, to block both K^+^ and Ca^++^ channels extensively. First, we examined the attenuating effect of EIBS in the paired experiment of high-K^+^ *vs* high-K^+^ with EIBS. One cochlea was incubated in the high-K^+^ (50 mM) extracellular solution (high-K^+^ group) and another cochlea was incubated with 50 mM K^+^ plus EIBS (K^+^+EIBS group). The ribbon numbers in both groups were normalized to that in the NES control group in Figure 1. Figure 4c shows that 95.5±3.0% of IHC ribbons in the K^+^+EIBS group were reserved for 50 mM K^+^ challenge (P=0.002, paired t test, two-tail). Figure 4d also shows that the ribbon volume in the high-K^+^+EIBS group was only increased 1.24±0.17 folds. In comparison with increase of 1.80±0.20 folds in the ribbon volume in the high-K^+^ group, the ribbon swelling in the high-K^+^+EIBS group was reduced by 70.0% (P=0.0007, paired t test, two-tail). We also examined the attenuating effect of EIBS in the paired experiment of NES *vs* K^+^+EIBS, in which one cochlea was incubated with NES and another cochlear was incubated in EIBS with 50 mM K^+^ (Supplementary Figure 2). In comparison with those in the control NES group, the ribbon number and volume in the K^+^+EIBS group were 98.9±3.96% and 1.23±0.21 folds, respectively (Supplementary Figure 2). There were no significant differences in the ribbon number and volume between the NES and K^+^+EIBS groups (P=0.73 and 0.32, respectively, paired t test, two-tail). In both paired experiments, the ribbon survival in the K^+^+EIBS group was >95% and the ribbon volume was also only increased ∼1.2 folds, similar to the volume increase caused by incubation procedure itself (Figure 1f). Thus, blockage of both Ca^++^ and K^+^ channels could almost completely attenuate high-K^+^ induced IHC ribbon degeneration and swelling.

Figure 5 shows the comparison of attenuations of K^+^ and Ca^++^ channel blockers on the ribbon degeneration and swelling. All of K^+^ and Ca^++^ blocker TEA, IBT, Co^++^, and EIBS could significantly attenuate high-K^+^ induced ribbon degeneration (P<0.001, one-way ANOVA with a Bonferroni correction, Figure 5a). EIBS had a maximum of attenuation and could allow 95.5% of ribbons survival for high-K^+^ challenge, whereas Co^++^ had 72.7% of minimum attenuation. There was significant difference between them (P=0.008, one-way ANOVA with a Bonferroni correction). However, there was no significant difference among attenuations of TEA, IBT, and Co^++^ in the ribbon degeneration (P=0.55, one-way ANOVA). For ribbon swelling (Figure 5b), there was no significant reduction after application of K^+^ channel blockers (P=1.00, one-way ANOVA). However, there was significant reduction in the ribbon volumes after application of Ca^++^ channel blockers. The ribbon volumes in the K^+^+Co^++^ group and the K^+^+EIBS group were only increased to 1.29±0.14 and 1.24±0.17 folds, respectively. In comparison with ∼1.8 folds of increase in the high-K^+^, K^+^-IBT, and K^+^-TEA groups, the ribbon swelling in the K^+^+Co^++^ group and K^+^+EIBS group were significantly reduced (P=0.0002, one-way ANOVA with a Bonferroni correction). However, there was no significant difference in the reduction of ribbon swelling between sole Co^++^ application and EIBS application (P=0.67, one-way ANOVA).

**Fig 5.**
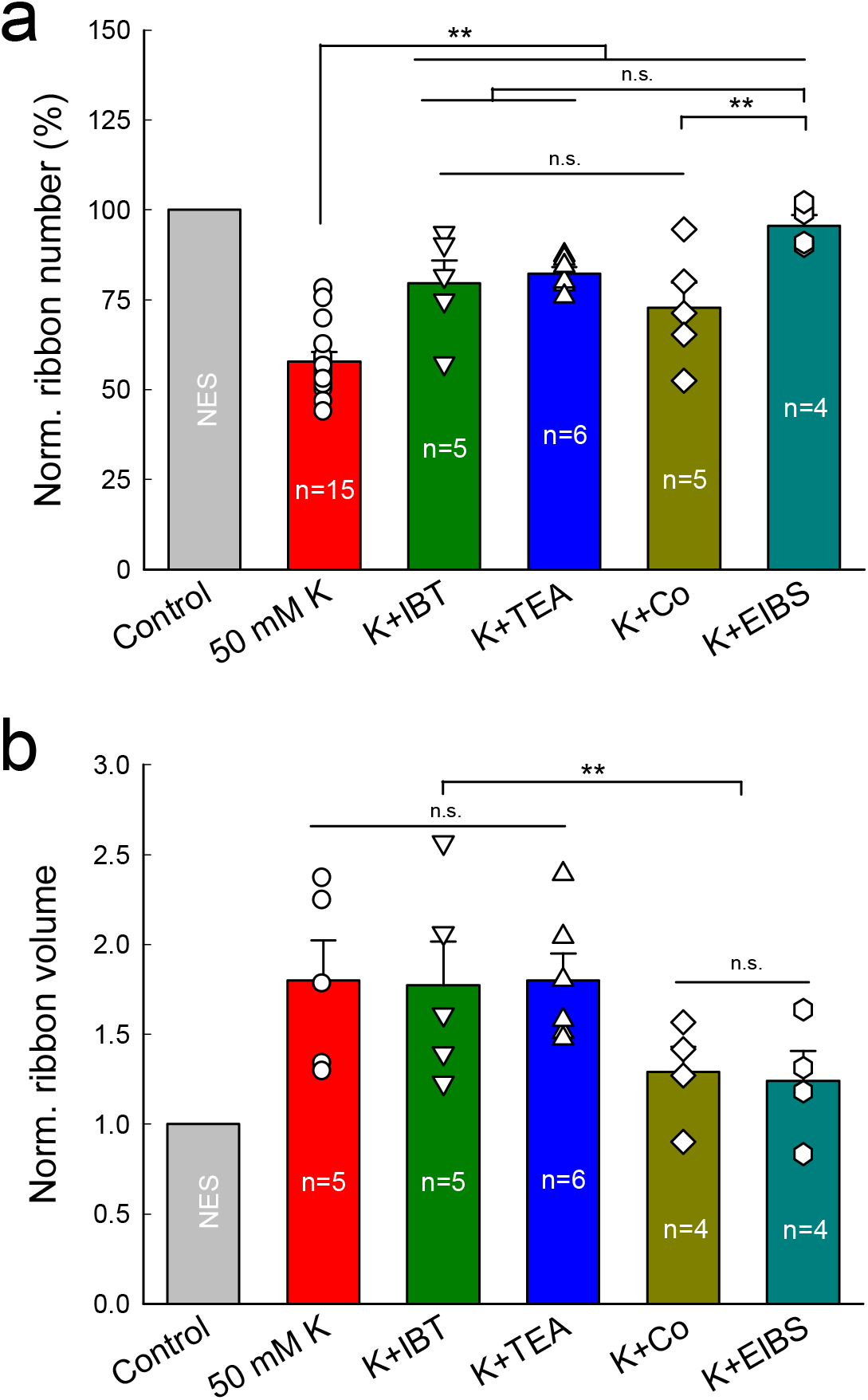
Comparison of attenuations of blockage of K^+^ and Ca^++^ channels on K^+^-induced IHC ribbon degeneration and swelling. Ribbon numbers and volumes in the K^+^ (50 mM) group, K^+^+IBT (0.1 μM) group, K^+^+TEA (20 mM) group, K^+^+Co^++^ (2 mM) group, and K^+^+EIBS group were normalized to those in the NES control group. N is the number of mice. **a:** Both K^+^ and Ca^++^ channel blockers attenuate ribbon degeneration. **b:** Ca^++^ channel blocker but not K^+^ channel blockers attenuates ribbon swelling. n.s.: no significance (P>0.05); **: P < 0.01, one-way ANOVA with a Bonferroni correction.

### GluR agonists cause IHC ribbon synapse swelling but not degeneration

To assess the effect of glutamate excitotoxicity on the IHC ribbon synapse degeneration, we investigated the effect of GluR agonists on IHC ribbon degeneration. In the same way, two cochleas in the same mouse were separately incubated with 0.2 mM glutamate and NES for 2 hr. Figure 6a shows that in comparison with those in the control NES group, the numbers of ribbons, GluRs, and synapses in the glutamate incubation group were 107±6.98, 104±6.04, and 93.5±5.75% (P=0.42, 0.42, and 0.30, paired t test, two-tail), respectively. Also, in comparison with those in the NES control group, the numbers of ribbons in the apical, middle, and basal turns in the glutamate group were 98.1±8.67, 101±6.43, and 110±10.3%, respectively (Figure 6b). There were no significant differences in ribbon numbers in the apical, middle, and basal turns between two groups (P=0.78, 0.91, and 0.45, paired t test, two-tail). However, glutamate could cause ribbon swelling (Figure 6c&d). In comparison with that in the control group, the volume of ribbons in the glutamate group was significantly increased 2.08±0.15 folds (Figure 6c, P=0.0002, paired t test, two-tail). The swelling was not significantly different from that caused by the high-K^+^ challenge (P=0.72, one-way ANOVA, Supplementary Figure 3b)

**Fig 6.**
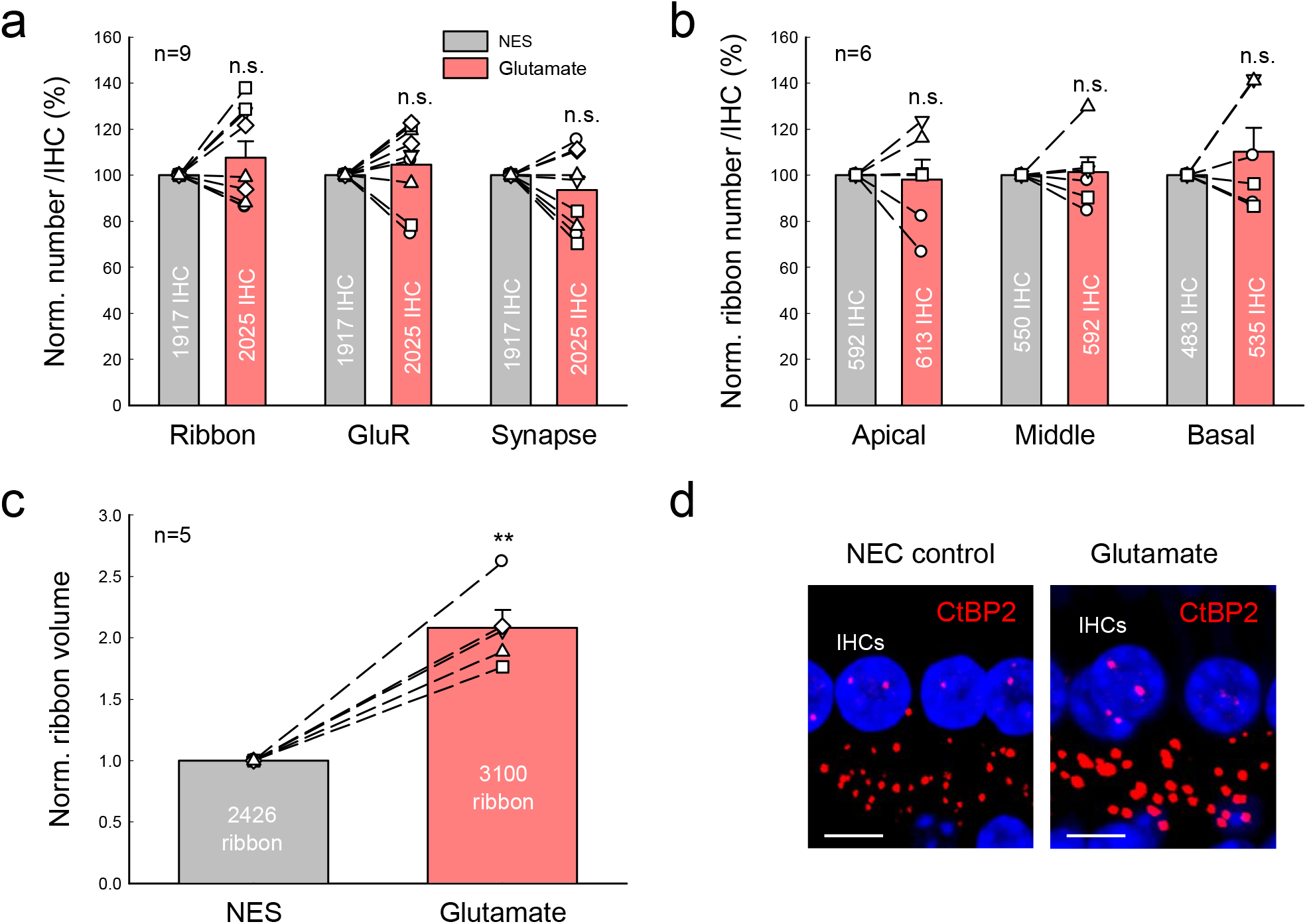
Glutamate causes ribbon swelling but not synapse degeneration. One cochlea was incubated in NES with 0.2 mM glutamate for 2 hr and another cochlea of the same mouse was simultaneously incubated with NES as control. Dashed lines connected two same symbols represent data collected from two cochleas of the same animal. N is the number of mice. **a:** Glutamate has no significant effect on IHC ribbon, GluR, and synapse degeneration. The numbers of ribbon, GluR, and synapse per IHC were pooled from all cochlear turns and averaged and normalized to those in the control cochlea of the same mouse. P=0.30-0.42, paired t test, two-tail. **b:** There are no significant ribbon degenerations in the different cochlear turns for glutamate challenge. The ribbon numbers per IHC in the different cochlear turns were normalized to those in the control cochlea of the same mouse. P=0.45-0.91, paired t test, two-tail. **c-d:** Glutamate-induced IHC ribbon swelling. Panel **d** is the immunofluorescence staining for CtBP2 at the middle cochlear turn area in the glutamate-treated group and control group. IHC ribbons in the glutamate treated cochlea appear enlarged. Scale bar: 10 µm. **: P < 0.01, paired t test, two-tail.

Application of GluR agonist Kainic acid (KA, 0.3 mM) or NMDA (0.3 mM) also did not cause ribbon degeneration (Figure 7a&c). In comparison with those in each NES control group, the ribbon numbers in KA and NMDA groups were 97.4±8.56 and 101.4±10.2%, respectively (P=0.70 and 0.83, paired t test, two-tail). However, like glutamate, KA, and NMDA caused ribbon swelling (Figure 7b&d). In comparison with that in the NES control group, the ribbon volumes in the KA group and NMDA group were significantly increased 1.65±0.20 and 1.75±0.24 folds, respectively (P=0.01 and 0.04, respectively, paired t test, two-tail). Thus, GluR agonists glutamate, KA, and NMDA could cause IHC ribbon swelling but not degeneration.

**Fig 7.**
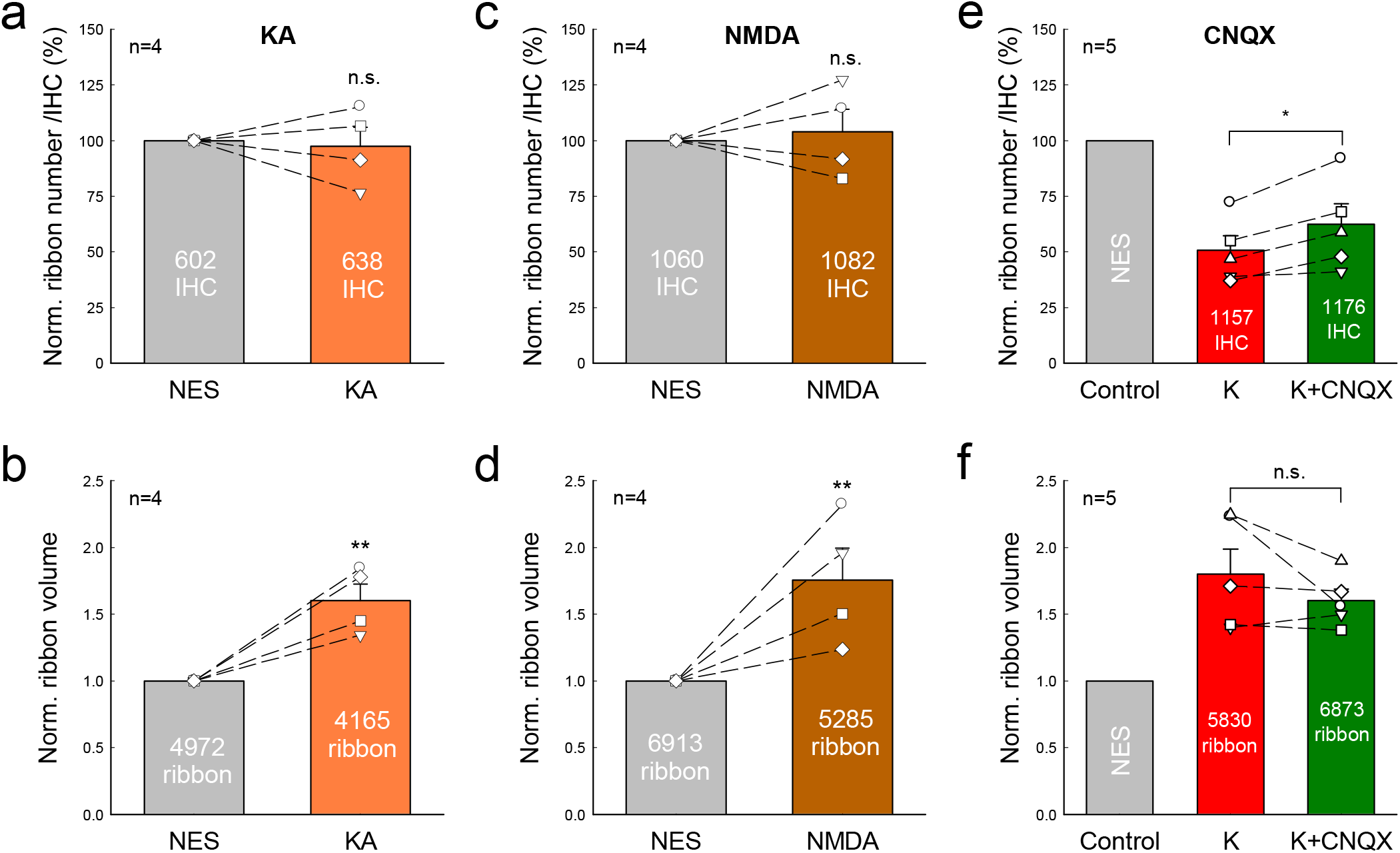
Effects of GluR agonist and antagonist on IHC ribbon degeneration and swelling. **a-d:** Application of 0.3 mM kainic acid (KA) or 0.3 mM NMDA causes ribbon swelling but not degeneration. The ribbon number and volume pooled from all cochlear turns were averaged and normalized to those in the control cochlea with NES incubation in the same mouse. N is the number of mice. **e-f:** Effects of a GluR antagonist CNQX on K^+^-induced ribbon degeneration and swelling. One cochlea was incubated in 50 mM K^+^ extracellular solution with 0.1 mM CNQX (K^+^+CNQX group) and another cochlea of the same mouse was incubated in 50 mM K^+^ extracellular solution (high-K^+^ group). Ribbon numbers and volumes in the high-K^+^ group and the K^+^+CNQX group were normalized to those in the NES control group in Figure 1. N is the number of mice. *: P<0.05, **: P < 0.01, and n.s.: no significance (P>0.05), paired t test, two-tail for all panels.

However, a GluR (AMPA/Kainate receptor) antagonist cyanquixaline (CNQX) could attenuate the K^+^-induced ribbon degeneration but not swelling (Figure 7e&f). After application of 0.1 mM CNQX with 50 mM K^+^, the survival of ribbon numbers had a small increase from 50.8±6.55% in the high-K^+^ group to 62.5±9.05% in the K^+^+CNQX group (P=0.014, paired t test, two-tail). However, there was no significant difference in the ribbon volume between the high-K^+^ group (1.80±0.18 folds) and the K^+^+CNQX group (1.60±0.09 folds) (P=0.22, paired t test, two-tail).

### IHC ribbon synapse degeneration after noise exposure

We further assessed IHC ribbon degeneration after noise exposure (Figure 8). We first assessed mouse hearing functional changes for noise exposure by auditory brainstem response (ABR) recording. Before noise exposure, ABR thresholds were 22.0±0.5, 22.3±1.08, 14.5±1.38, 19.5±0.82, 25.3±0.58, and 28.3±1.06 dB SPL in the control group and 21.3±0.67, 19.3±2.04, 11.0±1.45, 16.0±1.55, 22.0±1.04, and 27.0±2.41 dB SPL in the noise exposure group for click, 8, 16, 24, 32, and 40 kHz tone stimulations, respectively (Figure 8a). There were no significant differences in the pre-noise exposure ABR thresholds between two groups (P>0.05, one-way ANOVA). After exposure to white noise (95-98 dB SPL) for 2 hr, ABR thresholds measured at immediately stop of the exposure were 59.0±1.83, 52.7±3.40, 61.0±2.17, 67.3±1.95, 67.3±3.75, and 67.5±4.35 dB SPL for click, 8, 16, 24, 32, and 40 kHz tone-bursts, respectively (Figure 8a). In comparison with those at pre-noise exposure, the ABR thresholds were significantly elevated by ∼ 40 to 50 dB SPL (P< 0.001, one-way ANOVA with a Bonferroni correction). IHC ribbons in the control mice and noise-exposure mice were 13.9±0.34 and 8.97±0.62 per IHC, respectively (Figure 8b&c), and were reduced by 64.5% in the noise-exposure mice (P<0.001, one-way ANOVA with a Bonferroni correction). After noise exposure, ribbons also appeared swollen (Figure 8d). The ribbon volumes in the control mice and the noise-exposed mice were 0.418±0.029 and 0.647±0.085 μm^3^ (P= 0.035, one-way ANOVA with a Bonferroni correction), respectively; there was 1.55 folds increase in the noise exposure group.

**Fig 8.**
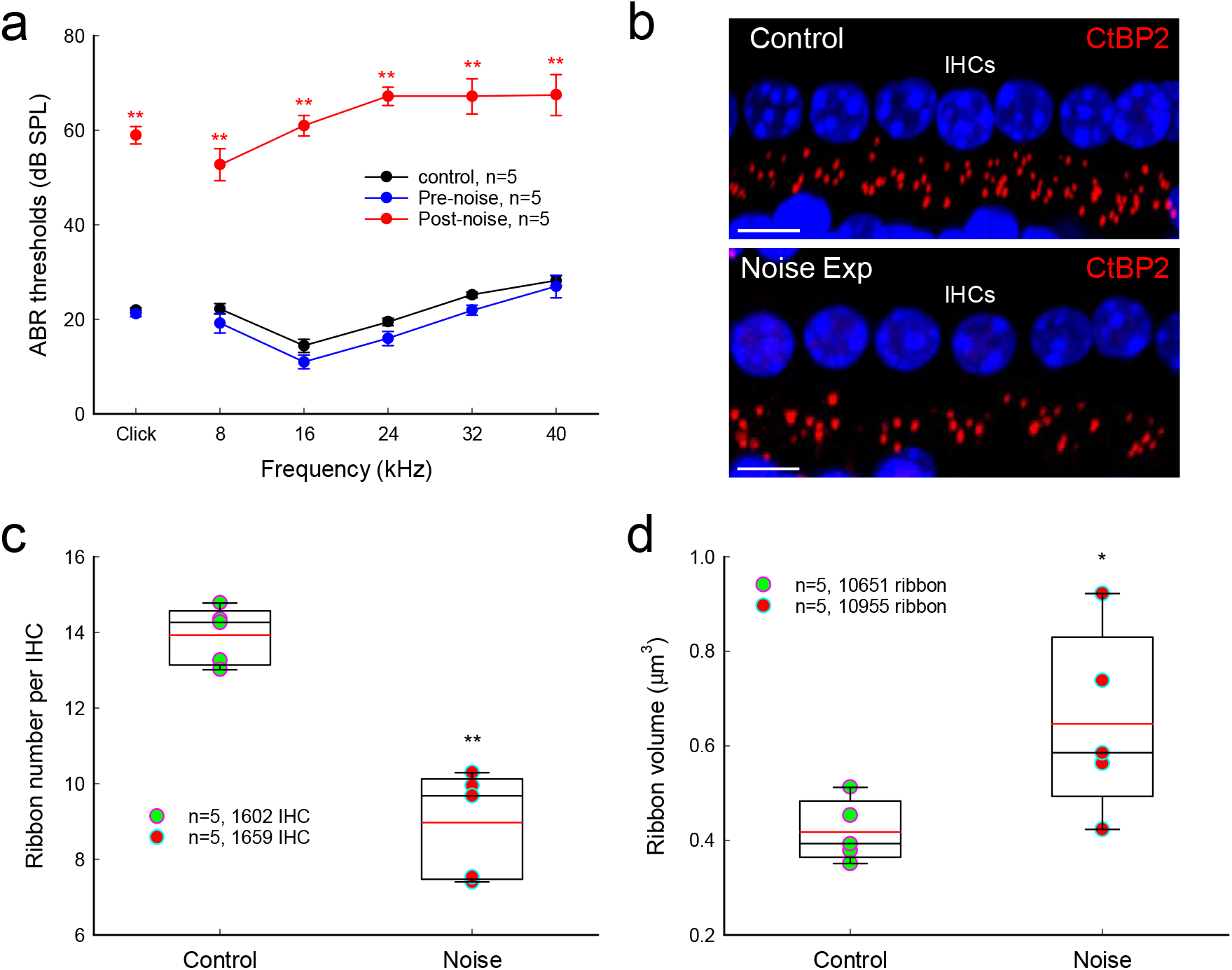
ABR threshold elevation and ribbon synapse degeneration after noise exposure. Mice were exposed to 95-98 dB SPL white-noise for 2 hr. After stop of noise exposure, ABR thresholds were immediately recorded and the inner ears were isolated after recording and fixed for immunofluorescence staining. **a:** Noise-induced ABR threshold elevation. After noise exposure, the ABR thresholds in the noise exposure mice were significantly increased about 45-55 dB SPL. N is the mouse number. **b:** Immunofluorescent staining for CtBP2 at IHC area at the middle turn in noise exposure and control mice. Scale bar: 10 µm. **c:** IHC ribbon synapse degeneration after noise exposure. Red lines in the boxes represent the mean level. N is the mouse number. **d:** IHC ribbon swelling after noise exposure. Red lines in the boxes represent the mean level. *: P < 0.05; **: P < 0.01, one-way ANOVA with a Bonferroni correction for all panels.

### No outer hair cell ribbon synapse degeneration for high-K^+^ challenge and noise exposure

However, there were no significant reductions in outer hair cell (OHC) ribbons for high-K^+^ challenges and noise exposure. Figure 9a-c show that there was no significant reduction in OHC ribbons after application of 50 mM K^+^. The numbers of OHC ribbons in the high-K^+^ (50 mM) group and the NES control group were 2.43±0.04 and 2.40±0.04 per OHC, respectively (Figure 9c, P= 0.31, one-way ANOVA). Also, there was no significant reduction in OHC ribbons in noise exposed mice (Figure 9d-f). The ribbon numbers in the noise exposure and control mice were 2.47±0.04 and 2.38±0.05 per OHC, respectively (Figure 9f, P=0.88, one-way ANOVA).

**Fig 9.**
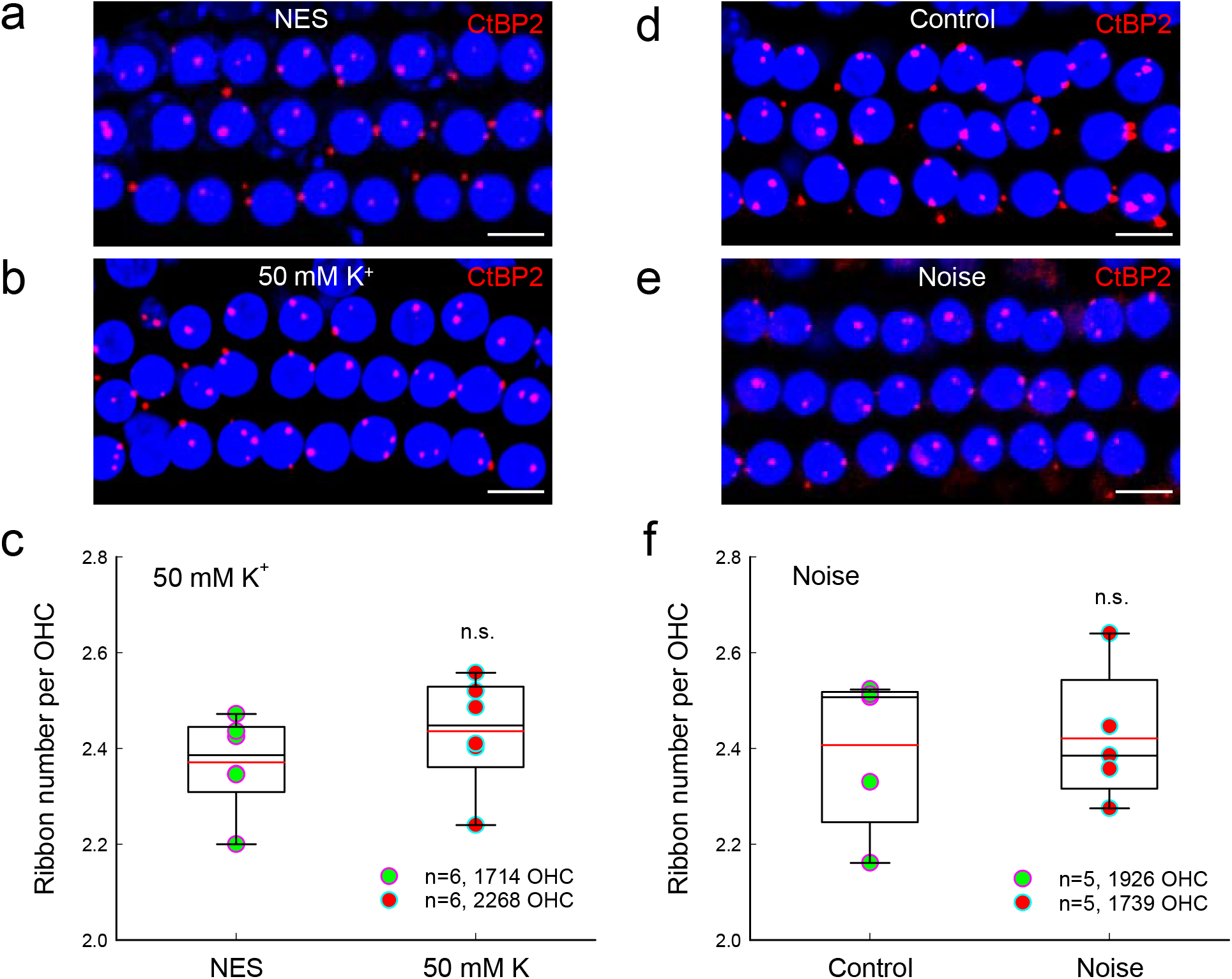
Outer hair cell ribbons have no degeneration for high K^+^ challenge and noise exposure. **a-b:** Immunofluorescent staining for CtBP2 in outer hair cells (OHCs) at the middle turn in the inner ears after incubation with 50 mM K^+^ extracellular solution and NES. Scale bar: 10 µm. **c:** OHC ribbons have no reduction after high-K^+^ (50 mM) challenge. There is no significant difference in OHC ribbon synapses between the high-K^+^ group and the control group (n.s.: no significance, P=0.31, one-way ANOVA). Data were averaged from all cochlear turns. N is the animal number. Red lines in the box represent the mean level. **d-e:** Immunofluorescent staining for CtBP2 in the OHC area at the middle cochlear turn in the noise exposure mice and control mice. Scale bar: 10 µm. **f:** OHC ribbons after noise exposure have no significant degeneration. Red lines in the box represent the mean level. There is no significant difference in OHC ribbon synapses between noise exposure mice and control mice. n.s.: no significance, P=0.88, one-way ANOVA.

## Discussion

In the present study, we found that elevation of extracellular K^+^ could cause IHC ribbon synapse degeneration and swelling (Figure 1). The degeneration was dose-dependent and increased as K^+^ concentration was increased (Figure 2). K^+^ channel blockers, especially BK channel blockers, could attenuate ribbon degeneration but not swelling, whereas the Ca^++^ channel blocker Co^++^ attenuated both ribbon degeneration and swelling (Figures 3-5). However, GluR agonists only caused ribbon swelling but not degeneration (Figures 6 and 7). A GluR (AMPA/Kainate receptor) antagonist CNQX could attenuate K^+^-induced ribbon degeneration but not swelling (Figure 7e&f). Finally, both high-K^+^ challenge and noise exposure caused only IHC ribbon degeneration but not OHC ribbon degeneration (Figures 8 and 9). These data suggest that high extracellular K^+^ can degenerate IHC ribbon synapse degeneration in HHL.

As mentioned above, the ribbon synapse degeneration has been hypothesized to result from glutamate excitotoxicity; acoustic overstimulation (noise exposure) results in excessive neurotransmitter glutamate release, thereby leading to excitotoxicity and hair cell and ribbon synapse degeneration (*17)*. However, cochlear synaptopathy in HHL has only ribbon synapse degeneration without hair cell death. Previous experiments also demonstrated that the presynaptic and postsynaptic components remained intact after perfusion of GluR agonists (*19,20*). In the present study, we found that application of GluR agonists caused ribbon swelling but not degeneration; there were no significant degenerations in presynaptic ribbons, postsynaptic GluRs, and synapses after application of GluR agonists (Figures 6 and 7, and Supplementary Figure 3). In addition, noise still could cause IHC ribbon swelling and degeneration in mice lacking vesicular glutamate transporter-3 and glutamate release (*18*). These results suggest that glutamate excitotoxicity may not be the main mechanism for IHC ribbon synapse degeneration.

When sensory cells and neurons are excited, intracellular K^+^ is released to the extracellular space to restore polarization. Excessive excitations of hair cells and neurons can increase K^+^ release and elevate extracellular K^+^, thereby leading to K^+^ excitotoxicity. In this study, we found that elevation of K^+^ caused IHC ribbon synapse degeneration (Figure 1 and Supplementary Figure 3). Blockage of K^+^ channels, in particular, BK channels attenuated this degeneration (Figures 3-5). The BK channel is a predominant type of K^+^ channels in mature IHCs (*25, 26, 29*). There are two types of BK channels in the mature IHCs (*27-29*): One can be blocked by IBT but is insensitive to Ca^++^ (*28, 30, 31*); another is IBT-insensitive but can be blocked by TEA and Cs^+^ (*28*). Both IBT and TEA could attenuate IHC ribbon degeneration (Figures 3 and 5). The TEA-sensitive BK channel is Ca^++^-dependent and can be activated by Ca^++^ flowing through L-type Ca^++^ channels (Cav1.3) (*28, 32*). In this study, application of Co^++^ attenuated high-K^+^ induced ribbon degeneration (Figures 4 and 5). High-K^+^ could depolarize IHCs to open Ca^++^ channels (*31-33*). Co^++^ may block Ca^++^ channels and consequently Ca^++^-dependent BK channels in IHCs. Moreover, it was reported that deficiency of BK channel activity by genetic deletion could increase resistance to noise-induced hearing loss in mice (*26*), although it had a report that BKα gene deletion could increase noise-induced hearing loss in low frequency regions with a higher noise vulnerability of OHCs (*34*). Taken together, these data suggest that BK channels may play a critical role in the IHC ribbon synapse degeneration. These data also suggest that BK channels may be a promising target for prevention and treatment of ribbon synapse degeneration in HHL.

In the experiments, we found that the GluR (AMPA/Kainate receptor) antagonist CNQX could attenuate high-K^+^ induced ribbon degeneration in IHCs (Figure 7e). IHCs have Ca^++^-permeable GluR4-assembled AMPA receptor expression (*35*). Thus, CNQX may block presynaptic GluR4 AMPA receptors and reduce Ca^++^ influx, thereby blocking Ca^++^-activated BK channels in IHCs to attenuate ribbon degeneration as Co^++^ did. Indeed, a recent study reported that blockage of Ca^++^-permeable AMPA receptors by IEM-1460 *in vivo* could attenuate noise-induced HHL and synaptopathy in mice (*36*). Noise exposure could increase extracellular K^+^ in the cochlea *in vivo*. Thus, blockage of Ca^++^-permeable AMPA receptors may consequently block Ca^++^-dependent BK channels as well in the IHCs to attenuate cochlear synaptopathy (Figure 7e&f), thereby eventually attenuating noise-induced HHL.

The present study demonstrated that TEA and IBT only attenuated ribbon degeneration (Figures 3 and 5), whereas Co^++^ could attenuate both ribbon degeneration and swelling (Figures 4 and 5). These data suggest that ribbon degeneration and swelling may underlie different mechanisms. Previous studies demonstrated that Ca^++^ excitotoxicity plays an important role in ribbon swelling (*20, 32, 36-37*). In the IHC, Ca^++^ can influx through Cav1.3 channels. High-K^+^ depolarizes the IHC and can open Cav1.3 channels to influx Ca^++^ (*29, 31-33*). Co^++^ could block Ca^++^ channels and eliminate such Ca^++^ influx, therefore attenuating ribbon swelling (Figures 4 and 5). Ca^++^ can also influx into IHCs through GluRs. As mentioned above, Ca^++^-permeable GluR4 AMPA receptors are expressed at presynaptic IHCs (*35*). Thus, application of GluR agonists could induce Ca^++^ influx and ribbon swelling but not degeneration (Figures 6 and 7) (*20, 38-40*). Since noise still caused IHC ribbon swelling in mice lacking the vesicular glutamate transporter-3 (*18*), both Ca^++^ channel-dependent and GluR-dependent pathways may work together *in vivo*. Currently, it is still unclear and under debating whether ribbon swelling is recoverable or eventually leads to degeneration (*18, 20, 37*). However, during two-hour incubation with GluR agonists in this study, we only found significant IHC ribbon swelling but not degeneration (Figures 6 and 7, and Supplementary Figure 3). Therefore, these data suggest that Ca^++^ excitotoxicity plays an important role in IHC ribbon swelling and may also have a role in the IHC ribbon degeneration *in vivo*.

IHC synapses *in vivo* are embedded in the low-K^+^ perilymph and K^+^-level in the perilymph is normally as low (∼ 5 mM). However, when the sensory cells and neurons are excited, they will release intracellular K^+^ to the perilymph increasing K^+^ concentration in the perilymph. The K^+^ concentration in the local extracellular space around hair cell’s synapse area even could be higher. Moreover, noise, aging or other factors could impair the integrity of the reticular lamina and the barrier between the endolymph and perilymph and allow K^+^-rich endolymph (K^+^ =∼150 mM) entering the perilymph (*21, 22*). All of these changes could lead to elevation of extracellular K^+^ concentration, thus causing K^+^ excitotoxicity and cell depolarization. In the brain, it is well-known that neuron damage by stroke can cause intracellular K^+^ release into the extracellular space leading to K^+^ excitotoxicity, which can cause the secondary cell death in the neighboring non-stroke area (*23, 24*). In this study, we found that high-extracellular K^+^ could cause IHC ribbon synapse degeneration (Figures 1-5). Like intensity dependence of ribbon synapse degeneration in noise exposure (*41*), the degeneration was dose-dependent (Figure 2). Blockage of K^+^ channels attenuated ribbon degeneration but not swelling (Figures 3-5). In addition, both K^+^ and noise caused IHC ribbon degeneration but not OHC ribbon degeneration (Figure 9). Therefore, these data suggest that K^+^ excitotoxicity may play an important role in the noise-induced cochlear synapse degeneration and HHL.

This study may also provide important information for mechanisms underlying other cochlear and hearing disorders, such as Meniere’s disease. Its symptoms have been considered to result from K^+^ intoxicating the hair cells and nerve fibers in the perilymph (*42, 43*). It was reported that there was a significant decrease in the number of afferent nerve endings and afferent synapses at the base of IHCs in Meniere’s disease (*44*), consistent with our finding in this study that high-K^+^ could cause IHC ribbon synapse degeneration (Figures 1-5). Our study, therefore, may also provide important information for understanding mechanisms of Meniere’s disease.

## Materials and Methods

### Animal and solution preparations

Both genders of adult CBA/CaJ mice (8-12 weeks old, Stock No: 000654, The Jackson Lab) were used. Mice were housed in a quiet individual room in basement with regular 12/12 light/dark cycle. The background noise level in the mouse room at mouse hearing range (4-70 kHz) is <30 dB SPL. All procedures and experiments followed in the use of animals were approved by the University of Kentucky’s Animal Care & Use Committee (UK: 00902M2005) and conformed to the standards of the NIH Guidelines for the Care and Use of Laboratory Animals.

The normal extracellular solution (NES) was prepared by 142 NaCl, 5.37 KCl, 1.47 MgCl_2_, 2 CaCl_2_, 10 HEPES in mM. The high-K^+^ extracellular solution was prepared by replacement of NaCl with KCl in the NES (*45*). The high-glutamate solution was made by adding 0.2 mM L-glutamate (cat. #49621, Sigma-Aldrich Inc) into the NES. The extracellular ionic blocking solution (EIBS) was prepared by 100 NaCl, 20 tetraethylammonium (TEA), 20 CsCl, 2 CoCl_2_, 1.47 MgCl_2_, 2 CaCl_2_, 10 HEPES in mM (*45*). Other incubation solutions were prepared by description in the text. All solutions were adjusted to pH 7.2 and 300 mOsm. The osmolality was verified by a freezing pressure micro-osmometer (Model 3300, Advance Instrument). Chemicals were purchased from Sigma-Aldrich Inc. (St. Louis, MS, USA). NMDA, Kainic acid (KA), and CNQX (6-cyano-7-nitroquinoxaline-2,3-dione) were purchased from TOCRIS Bioscience (Cat. #0114, 0222, and 1045, respectively). Iberiotoxin was purchased from EMD Millipore (Cat. #401002).

### Cochlear preparation and high-K^+^ challenge

The mouse cochlea was freshly isolated and the temporal bone was removed. The otic capsule was opened. The round window membrane was broken by a needle and a small hole was also made at the apical tip of the cochlea. The left or right cochlea was randomly selected and incubated in the high-K^+^ solution or other solutions; another cochlea in the same mouse was simultaneously incubated with NES or other control solutions for 2 hr, as the same time as noise exposure. At the beginning of incubation, gentle suctions and blows were performed several times on the round window and the apical hole to ensure the incubation solution quickly entering the cochlea. The complete exchange of perilymph could be verified with solution fluxed out of the round window or the apical hole by eyes under a stereo microscope (SMZ745, Nikon). After incubation for 2 hr, both the treated cochlea and the control cochlea were immediately fixed with 4% paraformaldehyde in PBS for 30 min for immunofluorescence staining.

### Preparation for histological examination

After fixation, the cochlea was dissected under a stereo microscope (SMZ745, Nikon) (*46, 47*). The bone that encloses the cochlea was gently crashed and removed. Then, the tectorial membrane and stria vascularis were carefully torn off by forceps. The basilar membrane with the organ of Corti (auditory sensory epithelium) was picked away by a sharpened needle and collected for staining. Starting from the cochlear apical region, the area at 25, 50, and 75% was considered as the apical, middle, and basal turn, respectively.

### Immunofluorescent staining and confocal microscopy

Both inner ears in the same mouse were stained in parallel to minimize variations of the performance. The isolated cochlear sensory epithelia were incubated in a blocking solution (10% goat serum and 1% BSA of PBS) with 0.1% Triton X-100 for 30 min. Then, the epithelia were incubated with primary antibodies for CtBP2 (1:200, mouse anti-CtBP2 IgG1, #612044, BD Bioscience) and GluR (1:500, mouse anti-GluR2 IgG2a, #MAB397, Millipore Corp.) in the blocking solution at 4^°^C overnight. In some of early experiments, only anti-CtBP2 antibody was used. After washout, the epithelia were incubated with corresponding second Alexa Fluor antibodies (1:500, goat anti-mouse IgG1 Alexa Fluor 568, A-21124, or with goat anti-mouse IgG2a Alexa Fluor 488, A-21131, Thermo Fisher Sci) at room temperature (23 ^°^C) for 2 hr. Before washout, 4’, 6-diamidino-2-phenylindole (DAPI, 50 μg/ml, D1306; Thermo Fisher Sci) was added to visualize the cell nuclei. After incubation for 2 min, the sensory epithelia were washed by PBS 3 times and whole-mounted on the glass-slide (*48*).

The stained epithelia were observed under a Nikon A1R confocal microscope system. Images were taken along the cochlear sensory epithelium from the apical turn to the basal turn. At each turn, at least 2-3 view-fields were imaged. At each field, serial sections were scanned along the Z-axis from the bottom to apical surface of the epithelium with a 0.25 μm step by use of a Nikon 100x (N/A: 1.45) Plan Apro oil objective at 0.25 μm/pixel resolution.

### Quantitative analysis and image presentation

The maximum intensity projection was generated from z-stack images at each image field. The numbers of ribbons (puncta of CtBP2 labeling), GluR labeling, and synapses (contacted or overlaped CtBP2 and GluR labeling) in the IHC region were separately accounted by NIS Elements AR Analysis software (Nikon). The IHC numbers in the field were also accounted. The number per IHC was calculated and averaged from all imaged fields in each cochlear turn or the whole cochlea. Then, they were normalized to those in the control ear of the same mouse. Finally, the numbers among different animals within a given experiment were averaged and compared. In some cases, the ribbon number per OHC was also calculated.

The ribbon volumes were measured from the 3D-image created from z-stack sections by Imaris software (Bitplane Inc., MA, USA). The vesicle volume measure function in the Imaris software was used with 0.15 μm smoothing. The volume measurement was performed by defining a minimum threshold above local background and a minimum acceptable Guissian distribution of voxel intensity to minimize false-positive noise. Similar parameters were used in the measurements among different mice. The volumes of IHC ribbons were measured from at least 3 different imaging fields in each cochlea and averaged. The means was normalized to that of the control ear of the same animal and further averaged among different animals within a given experiment (*49*).

### Noise exposure

Littermate mice (8-10 weeks old) in one cage were randomly separated to the noise exposure group and the control group to ensure age and sex-matched control. Mice were awake in a small cage under loud-speakers in a sound-proof chamber and exposed to white-noise (95-98 dB SPL) for 2 hr, one time. Noise was amplified by a power amplifier (CSA 240Z, JBL Commercial, UT, USA). Sound pressure level and spectrum of noise in the cage were measured and verified prior to placement of mice.

### Auditory brainstem response recording

Auditory brainstem response (ABR) was recorded by use of a Tucker-Davis R3 workstation with ES-1 high frequency speaker (Tucker-Davis Tech. Alachua, FL) (*46, 47*). For ABR recording, mice were anesthetized with a mixture of Ketamine and Xylazine (1 ml Ketamine+0.55 ml Xylazine+8.5 ml saline, 0.1 ml/10 g body weight, i.p.). Body temperature was maintained at 37–38°C by placing anesthetized mice on an isothermal pad (Deltaphase, model 39dp, Braintree Scientific Inc., Massachusetts). ABR was recorded by stimulation with clicks in alternative polarity and tone bursts (4-40 kHz) from 80 to 10 dB SPL in a 5 dB step in a double-walled sound isolation room. The signal was amplified (50,000x), filtered (300-3,000 Hz), and averaged by 500 times. The ABR threshold was determined by the lowest level at which an ABR can be recognized (*46, 47*). ABR was recorded at 1-2 day before noise exposure as reference and intermediately after stop of noise exposure to assess the threshold shift by noise exposure.

### Statistics and Reproducibility

Data were plotted by SigmaPlot (version 13, Systat Software) and statistically analyzed by Excel (Microsoft Office 360) or SPSS (version 25, SPSS Inc.; Chicago, IL). Error bars represent SEM. Parametric and nonparametric data comparisons were performed using one-way ANOVA or Student t tests after assessment of normality and variance. The paired, two-tailed Student’s t test was used for comparison of measures within the same group. The threshold for significance was p= 0.05. Bonferroni *post hoc* test was used in ANOVA.

For *in vitro* experiments, two inner ears in one mouse were used at each time. Each *in vitro* experiment was repeated at least 4 times to verify the reproducibility. Major experiments for high-K^+^ challenge and glutamate treatment were repeated for more than 10 times with different experimenters (see Supplementary Figure 3). For *in vivo* noise exposure experiment, only one or two mice were exposed at each time. Totally, 5 mice were exposed to noise and 3 times were repeated.

### Data Availability

All data needed to evaluate the conclusions in the paper are present in the paper and/or the Supplementary Materials. Additional data related to this paper may be requested from the authors.

## Acknowledgement

This work was supported by NIH R01 DC 017025 and R56 DC 016585 to HBZ.

## Author Contributions

HBZ conceived the general framework of this study. YZ, LML, and HBZ performed the experiments. HBZ, YZ, and LML analyzed data. HBZ wrote the paper. All authors reviewed the manuscript and provided the input.

---

**Supplementary Figure 1.**
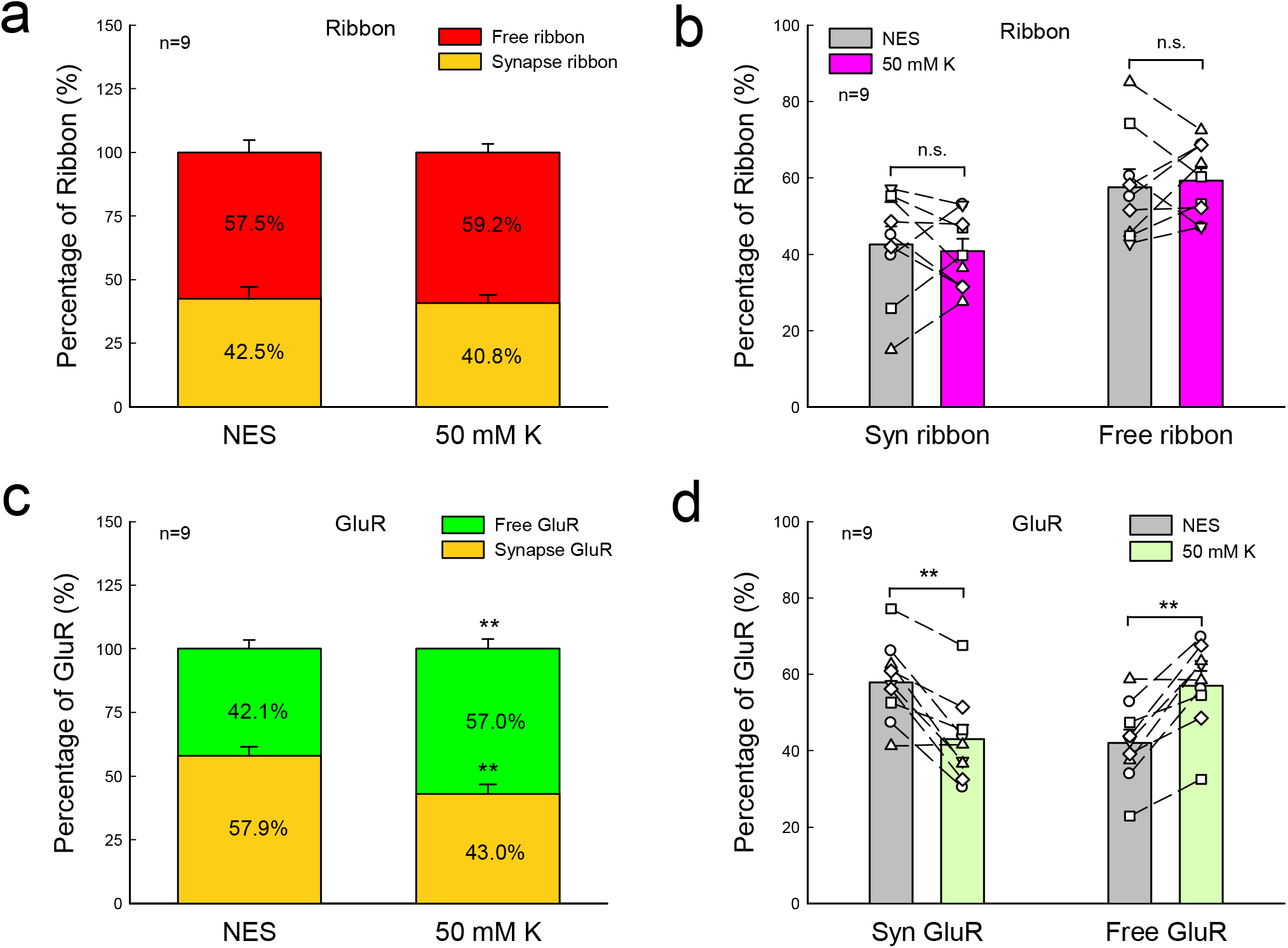
Changes in synaptic ribbons, free ribbons, synaptic GluRs, and free GluRs after high-K^+^ (50 mM) treatment. The percentages of synaptic ribbons and free ribbons in the NES group and high-K^+^ group were calculated by dividing total ribbon number in each group. The same calculation was also applied to GluRs. **a-d** : Changes in synaptic and free ribbons after high-K^+^ challenge. There are no significant changes in percentages of synaptic and free ribbons between the high-K^+^ group and the NES control group, indicating that K^+^ degenerated both synaptic and free ribbons. N is the animal number. P=0.68, paired t test, two-tail. **c-d** : Changes in synaptic and free GluRs after high-K^+^ challenge. The percentage of synaptic GluRs in the high-K^+^ group is significantly reduced, indicating major loss of synaptic GluRs after high-K^+^ challenge. **: P < 0.01, paired t test, two-tail.

**Supplementary Figure 2.**
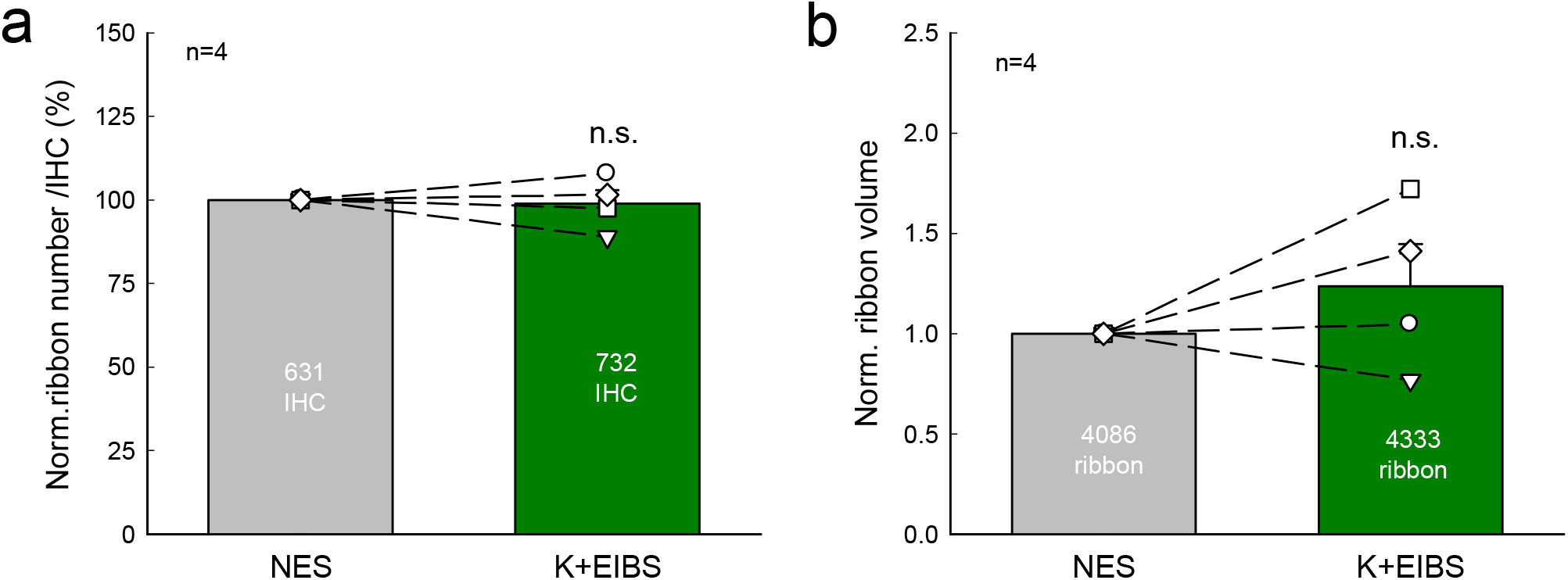
Elimination of K^+^-induced IHC ribbon degeneration and swelling by extracellular ionic blocking solution (EIBS) in the paired experiment with normal extracellular solution (NES) *vs* K^+^+EIBS. One cochlea was incubated in EIBS with 50 mM K^+^ (K^+^+EIBS) and another cochlea of the same mouse was incubated in the NES as control. The data were normalized to those in the control cochlea in the same mouse. N is the number of mice. **a** : IHC ribbon numbers in the K^+^+EIBS group have no significant reduction for 50 mM K^+^ challenge. There is no significant difference in IHC ribbon numbers between the K^+^+EIBS group and the NES control group. n.s.: no significance (P=0.52), paired t test, two-tail. **b** : IHC ribbon volumes in the K^+^+EIBS group are not significantly increased for 50 mM K^+^ challenge. There is no significant difference in ribbon volumes between the K^+^+EIBS group and the NES control group. n.s.: no significance (P=0.94), paired t test, two-tail.

**Supplementary Figure 3.**
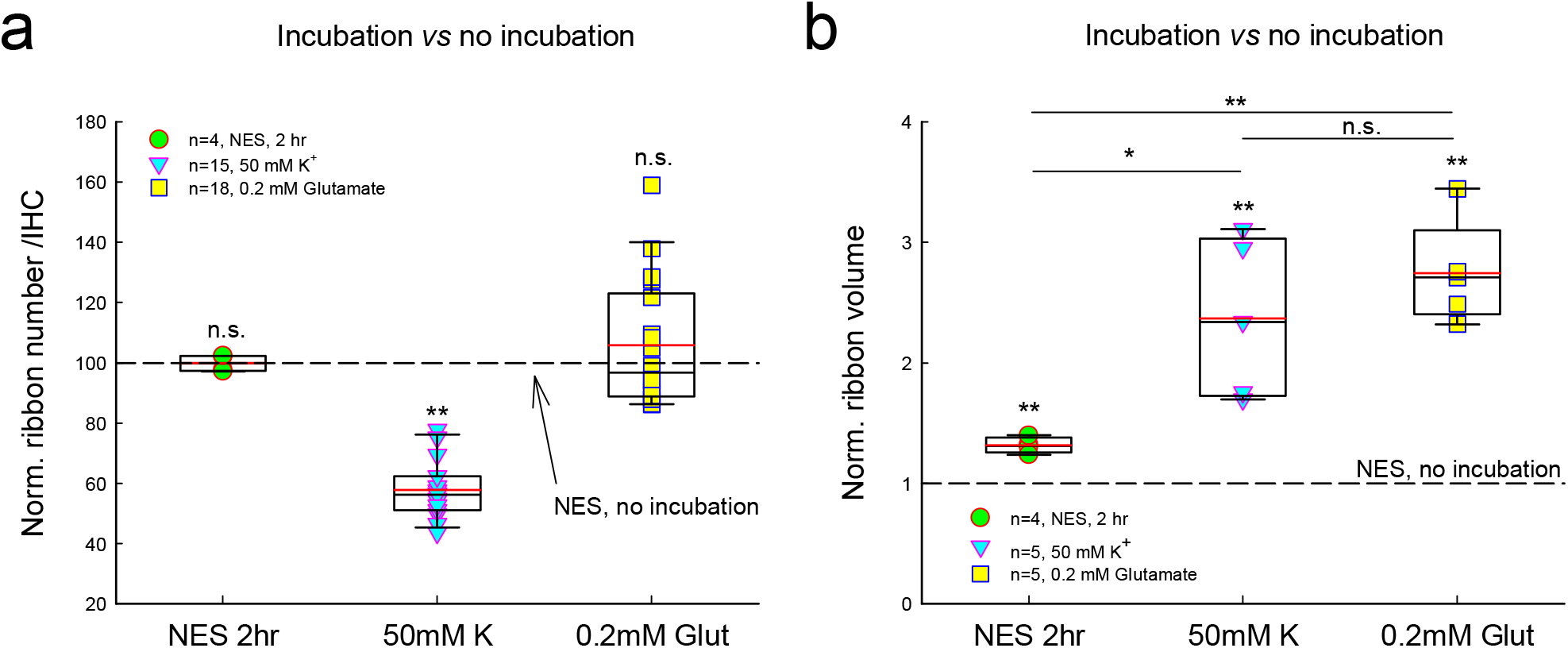
Comparisons of effects of incubation procedure, high-K^+^ (50 mM) challenge, and glutamate (0.2 mM) treatment on IHC ribbon number and volume. The ribbon number per IHC and ribbon volume were normalized to those in the non-incubation control group, which are indicated by dashed lines in the figures. Red lines in the box represent the mean level. N is the number of mice. **a :** The effect on ribbon numbers. In comparison with that in the non-incubation group (indicated by a dashed line), the normalized ribbon number per IHC after incubation with NES for 2 hr, 50 mM K^+^ challenge, and 0.2 mM glutamate application was 99.9±1.40, 57.8±2.54, and 105.9±4.95%, respectively. Application of 50 mM K^+^ caused significant reduction in IHC ribbons. However, incubation procedure and glutamate treatment had no significant effect on IHC ribbon degeneration. **: P<0.01, n.s.: no significance, P>0.05, one-way ANOVA with a Bonferroni correction. **b :** The effect on ribbon volumes. In comparison with that in the non-incubation group (indicated by a dashed line), the IHC ribbon volumes in the incubation with NES for 2 hr, 50 mM K^+^ challenge, and 0.2 mM glutamate treatment were significantly increased 1.32±0.03, 2.37±0.29, and 2.74±0.19 folds, respectively. The increases (2.37-2.74 folds) in 50 mM K^+^ or 0.2 mM glutamate treatment were significantly larger than that (1.32 folds) in the incubation with NES for 2 hr. **: P<0.01; *: P<0.05; n.s.: no significance, P>=0.05, one-way ANOVA with a Bonferroni correction.

